# Stable nebulization and muco-trapping properties of Regdanvimab/IN-006 supports its development as a potent, dose-saving inhaled therapy for COVID-19

**DOI:** 10.1101/2022.02.27.482162

**Authors:** Morgan McSweeney, Ian Stewart, Zach Richardson, Hyunah Kang, Yoona Park, Cheolmin Kim, Karthik Tiruthani, Whitney Wolf, Alison Schaefer, Priya Kumar, Harendra Aurora, Jeff Hutchins, Jong Moon Cho, Anthony J. Hickey, Soo Young Lee, Samuel Lai

**Author notes:** Corresponding Author: Samuel K. Lai, Ph.D., Division of Pharmacoengineering and Molecular Pharmaceutics, Eshelman School of Pharmacy, University of North Carolina-Chapel Hill, Chapel Hill, NC, 27599, USA.

## Abstract

The respiratory tract represents the key target for antiviral delivery in early interventions to prevent severe COVID-19. While neutralizing monoclonal antibodies (mAb) possess considerable efficacy, their current reliance on parenteral dosing necessitates very large doses and places a substantial burden on the healthcare system. In contrast, direct inhaled delivery of mAb therapeutics offers the convenience of self-dosing at home, as well as much more efficient mAb delivery to the respiratory tract. Here, building on our previous discovery of Fc-mucin interactions crosslinking viruses to mucins, we showed that regdanvimab, a potent neutralizing mAb already approved for COVID-19 in several countries around the world, can effectively trap SARS-CoV-2 virus-like-particles in fresh human airway mucus. IN-006, a reformulation of Regdanvimab, was stably nebulized across a wide range of concentrations, with no loss of activity and no formation of aggregates. Finally, nebulized delivery of IN-006 resulted in 100-fold greater mAb levels in the lungs of rats compared to serum, in marked contrast to intravenously dosed mAbs. These results not only support our current efforts to evaluate the safety and efficacy of IN-006 in clinical trials, but more broadly substantiate nebulized delivery of human antiviral mAbs as a new paradigm in treating SARS-CoV-2 and other respiratory pathologies.

## Introduction

Most viruses that cause acute respiratory infections (ARIs), including influenza [1–4], RSV [5–9], PIV [10] and the betacoronavirus HKU1 [11], infect almost exclusively via the apical (luminal) side of the airway epithelium, as revealed by studies using well-differentiated, polarized human airway epithelial (WD-HAE) cultures grown at the air-liquid interface [12]. In contrast, there is little-to-no productive infection when viruses are introduced into the basal (serosal) compartment in WD-HAE cultures. Importantly, infected cells appear to predominantly shed progeny viruses back into the apical compartment (i.e., into airway mucus (AM) secretions), with only rare shedding of virus into the basal compartment. This unique pathophysiology is shared by SARS-CoV-1, which only productively infects WD-HAE cultures when the virus is inoculated apically, with no appreciable infection when the same amount of virus is inoculated basally [13]. There is ^~^1,000-fold greater virus shed into the apical compartment relative to the basal compartment. The near exclusive apical infection and shedding of SARS-CoV-1 is consistent with the apical trafficking of ACE2 in airway biopsy tissues [14] and in WD-HAE cultures *in vitro* [13, 15, 16]. Not surprisingly, given that SARS-CoV-2 binds the same ACE2 receptor as SARS-CoV-1 for cellular entry, SARS-CoV-2 also undergoes the same preferential apical infection and shedding [13, 15, 16]. This pathophysiology is consistent with the substantial time window between initial appearance of upper respiratory tract symptoms and the development of pulmonary and systemic morbidities that necessitate hospitalization.

Given the concentration of SARS-CoV-2 in the respiratory tract, and the resulting respiratory tract symptoms and morbidities, direct delivery of potent neutralizing mAbs to the site of infection should have been preferred. Nevertheless, every antiviral mAb that has received full approval or EUA to date is dosed either by intravenous infusion or subcutaneous/intramuscular injections, despite prior studies showing only a small fraction of systemically dosed mAb reaches the respiratory tract in animal models [17–20] and in human studies [21]. The lack of efforts advancing inhaled delivery of mAb is likely due in part to prior work suggesting mAbs can aggregate and lose binding activity following nebulization [22, 23]. This problem is particularly evident with jet and ultrasonic nebulizers, where droplet recirculation, as well as heat and shear stresses, increase the aggregation of biomolecules and leave large residual quantities of unnebulized drug [24, 25]. Indeed, an earlier study delivering omalizumab using jet nebulizers for asthma was thought to possibly generate protein aggregates [26]. To date, there has been no report on stable nebulization of a fully human mAb that has been advanced through late-stage clinical trials. Nevertheless, a number of protein therapeutics have been stably nebulized using vibrating mesh nebulizers (VMN) as part of chronic treatment regimens [27–30]. This offers the potential that human mAbs, if appropriately formulated [22], can also be stably nebulized using VMNs, with no loss in binding and no aggregation.

Given the public health urgencies of the COVID-19 pandemic, we were motivated to advance an inhaled antiviral therapy using a full length, broadly neutralizing mAb. Regdanvimab is one of few anti-SARS-CoV-2 neutralizing mAbs that have received approval from either the FDA or EMA. Regdanvimab, administered by IV infusion at 40 mg/kg, provided a 72% reduction in risk of hospitalization and shortened the recovery time by ^~^5 days compared to placebo in its global Phase 3 clinical trial [31]. These results make regdanvimab a highly promising mAb for developing an inhaled COVID-19 therapy. Here, we report that IN-006, a reformulation of regdanvimab for nebulized delivery, can facilitate effective trapping of SARS-CoV-2 virus-like particles in human airway mucus (AM) and can be stably nebulized across a range of mAb concentrations with no loss of activity and no detectable aggregation. These unique properties of IN-006 enable us to achieve far higher mAb concentrations in the lung *in vivo*.

## Results

### IN-006 effectively traps SARS-CoV-2 VLPs in human airway mucus

To evaluate whether IN-006 can trap SARS-CoV-2 in human AM, we prepared fluorescent SARS-2 virus-like-particles (VLP) by co-expressing S protein with GAG-mCherry fusion construct and performed high resolution multiple particle tracking to quantify the mobility of hundreds to thousands of individual virions in each sample of fresh human AM isolated from extubated endotracheal tubes. In human AM treated with control mAb (motavizumab), SARS-CoV-2 VLPs exhibited rapid mobility that spanned microns within seconds (Figure 1A), as quantified by its ensemble averaged mean squared displacements (<MSD>) over time (Figure 1B). The average effective diffusivity (<D_eff_>) of SARS-CoV-2 VLPs was 0.23 μm^2^/s (Figure 1D), and rapid mobility of SARS-CoV-2 VLPs were observed in nearly all AM samples tested (Figure 1E, F). Such high diffusivities of SARS-CoV-2 VLPs are consistent with the high rates of SARS-CoV-2 transmission, and imply that immune-naïve AM does not pose an adequate diffusional barrier that effectively limits SARS-CoV-2 from reaching and infecting target epithelial cells along the respiratory tract.

**Figure 1:**
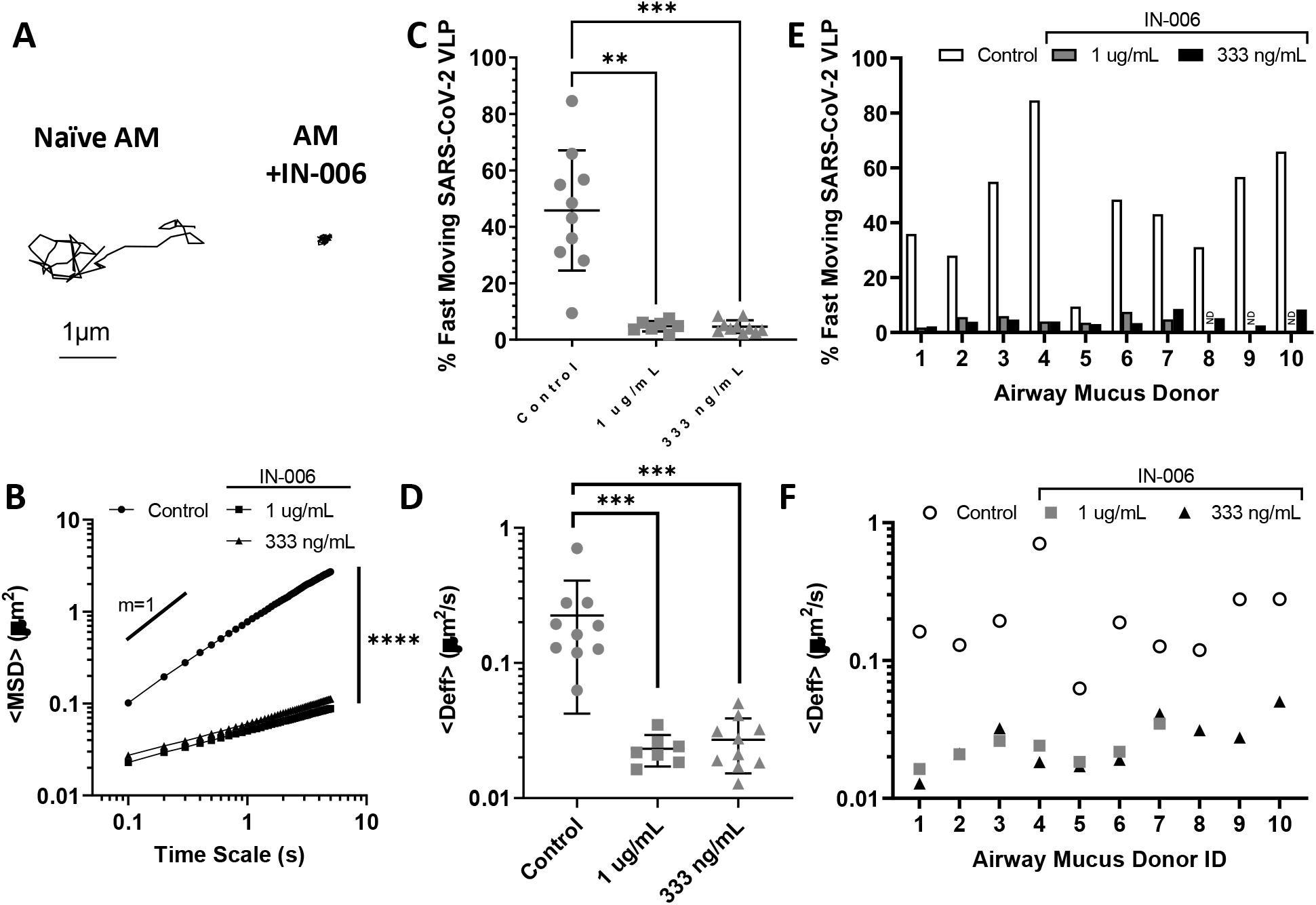
IN-006 effectively traps SARS-CoV-2 VLPs in fresh, undiluted human airway mucus (AM). **(A)** Representative traces of SARS-CoV-2 VLPs in AM treated with control mAb vs. in AM treated with IN-006. **(B)** Mean squared displacements of SARS-CoV-2 VLPs over time scales. **(C)** Fraction of rapidly diffusing SARS-CoV-2 VLPs and **(D)** effective diffusivities (<Deff>) of SARS-CoV-2 VLPs in AM treated with control mAb or IN-006 to a final concentration of 1 μg/mL and 333 ng/mL. A total of 10 independent donor samples were tested. **(E)** Fraction of rapidly diffusing SARS-CoV-2 VLPs and **(F)** effective diffusivities (<Deff>) of SARS-CoV-2 VLPs in each of the independent AM tested. Due to specimen volume limitations, we only assessed the muco-trapping potencies of IN-006 at 333 ng/mL for donor ID 8, 9 and 10 (ND = No Data). Repeated Measures One-Way Anova with post-hoc Dunnett’s test on log-transformed (diffusivity, MSD) or untransformed (% fast moving) data (*p<0.05, **p<0.01, ***p<0.001, ****p<0.0001).

We next assessed the mobility of SARS-CoV-2 in AM treated with IN-006 (to a final concentration of 1 μg/mL). In sharp contrast to treatment with control mAb, the mobility of SARS-CoV-2 VLPs were far more restrained in AM treated first with IN-006 (Figure 1A), as indicated by the much smaller <MSD> over time scale, and the reduced slope of the <MSD> vs. time scale plot (Figure 1B). IN-006 mediated trapping of SARS-CoV-2 was consistently observed in every AM sample tested, as reflected by the significant reduction in <Deff> in every AM specimens tested (Figure 1F). On average, compared to naïve AM, IN-006 added to AM to a final concentration of 1 μm/mL and 333 ng/mL reduced the <D_eff_> of SARS-CoV-2 VLPs by ^~^10-fold and ^~^8 fold, respectively; the <D_eff_> of virions in IN-006-treated AM were slowed ^~^190-and 160-fold in AM compared to their theoretical diffusivities in water. These results confirmed IN-006 effectively reduced the rapid mobility of SARS-CoV-2 in human AM.

Virions that possess the greatest diffusivity (i.e., the most mobile fractions), by definition, are more likely to diffuse across the mucus layer and infect the underlying epithelium before mucus is purged by natural clearance mechanisms. Thus, we sought to assess the effect of IN-006 in limiting the fraction of SARS-CoV-2 VLPs that could most readily penetrate across AM. We quantitatively defined the fast-moving subpopulation of SARS-CoV-2 as virions that possess sufficient mobility to penetrate through a physiologically thick AM layer (50 μm) in one hour, which yielded a minimum Deff ≥ 0.347 μm^2^/s). This fast-moving population was reduced from 46% in naïve AM to 4.8% and 4.6% in AM containing 1 μm/mL and 333 ng/mL of IN-006, respectively (Figure 1C), and was consistently observed in every AM specimens tested (Figure 1E). These results firmly underscore the effectiveness of muco-trapping IN-006 in limiting the mucus permeation of SARS-CoV-2 VLPs.

### IN-006 can be stably nebulized across a range of concentrations

The most efficient method to deliver mAb to the respiratory tract is by direct inhalation [20]; we thus tested whether we could generate IN-006-containing aerosols that are suitable for pulmonary deposition using a VMN. We first determined the aerodynamic particle size distribution (APSD) of the aerosols, as the resulting droplet sizes directly influence the site of deposition within the airways [32]. As a general guide, aerosols smaller than ^~^2.5 μm are preferentially deposited in the deep lung, between 2.5 to 5 μm preferentially in the lower airways, and aerosols ^~^5-10 μm in diameter preferentially in the upper airways, including nasopharyngeal and oropharyngeal regions [32]. Based on earlier unpublished work with nebulizing mAbs against RSV, we first utilized the Copley Scientific Next Generation Impactor (NGI) to measure the APSD of IN-006 nebulized at 20 and 30 mg/mL. The mass median aerodynamic diameter (MMAD) was 5.7 ± 0.08 μm and 5.3 ± 0.2 μm (Figure 2A), with a fine particle fraction (FPF; particulates <5 μm) of 47 ± 1% and 51 ± 1%, respectively. These results suggest that IN-006 could be nebulized in aerosols suitable for deposition throughout the entire respiratory tract.

**Figure 2:**
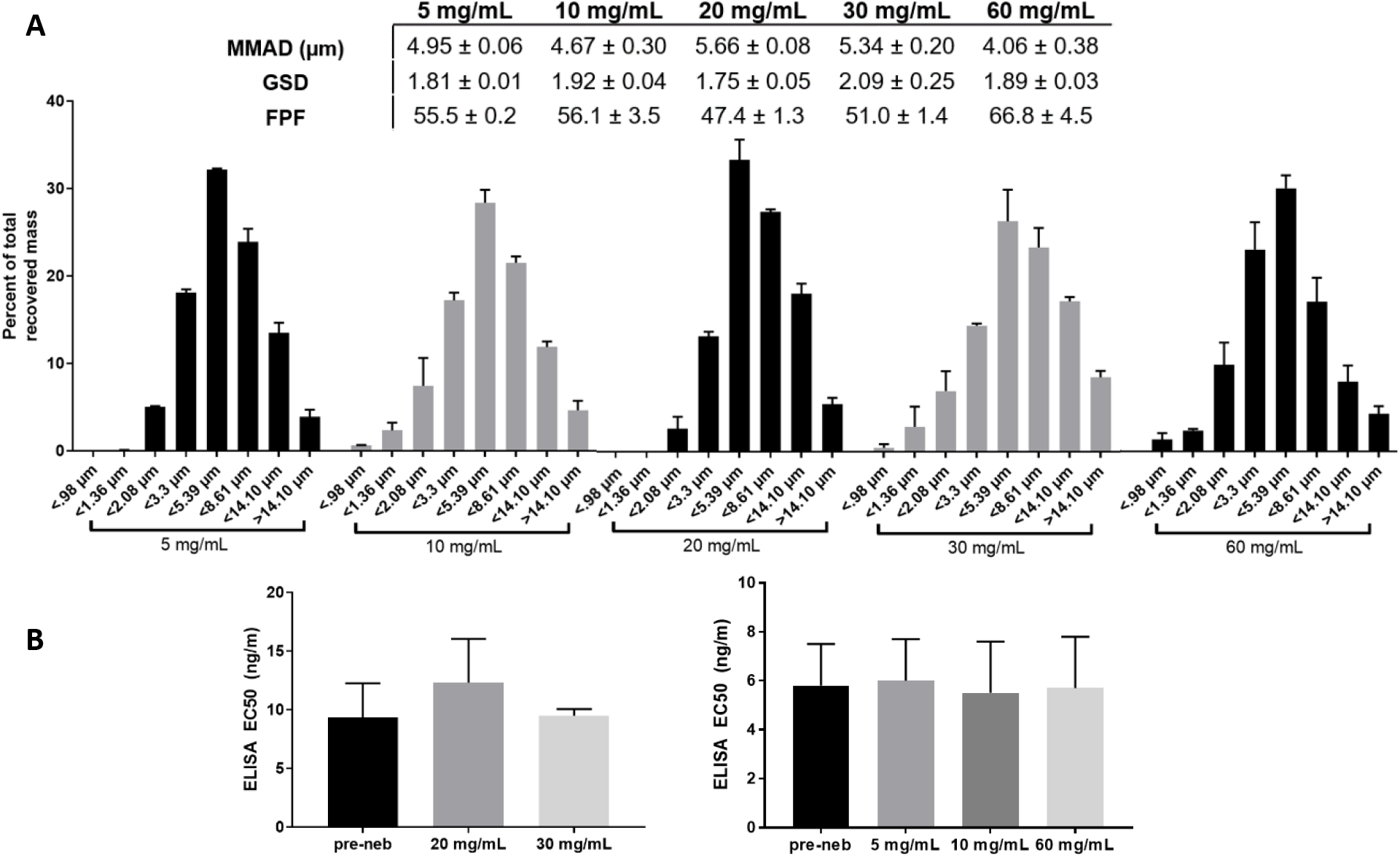
IN-006 retains stable binding activity after nebulization at various concentrations. IN-006 was formulated at 5, 10, 20, 30 or 60 mg/mL and nebulized using a Phillips InnoSpire Go VMN into an NGI at 15 L/min. **A)** Aerosol particle size distribution following nebulization at each concentration, plotted as a fraction of total dose recovered (100 × mass on NGI stage / sum of mass on all stages). Summary APSD characteristics of IN-006 nebulized at varying concentrations are shown in table. **B)** Binding activity of IN-006 pre- and post-nebulization, as determined by anti-SARS-CoV-2 S protein ELISA and calculation of EC_50_. Post-nebulization samples were collected from Stage 4 of the NGI (<5.39 μm). EC50 experiments for 20 mg/mL and 30 mg/mL formulations were conducted separately from those conducted subsequently with 5, 10, and 60 mg/mL, and are plotted separately.

To determine whether the binding affinity of IN-006 was preserved during nebulization, we collected nebulized IN-006 using a two-stage glass twin impinger setup in accordance with European Pharmacopoeia 2.9.18, and measured the binding affinity (EC_50_) of the recovered nebulized IN-006 via S-protein ELISA. In this impinger setup, aerosols with diameter greater than ^~^6.4 μm are primarily collected in the upper chamber, and particles smaller than 6.4 μm are primarily collected in the lower chamber. For IN-006 nebulized at 20 mg/mL, the EC_50_ of pre-nebulized IN-006, as well as IN-006 collected in the upper and lower impinger chambers were measured to be 9 ± 4 ng/mL, 12 ± 4 ng/mL, and 10 ± 1 ng/mL, respectively (Figure 2B). Likewise, for IN-006 nebulized at 30 mg/mL, the EC_50_ for pre-nebulized IN-006 and IN-006 recovered from the upper and lower impinger chambers were 10 ± 2 ng/mL, 10 ± 2, and 10 ± 1 ng/mL, respectively. To determine if mAb aggregates were generated by nebulization, we performed dynamic light scattering (DLS) on the recovered nebulized IN-006. IN-006 recovered from both the upper and lower impinger chambers exhibited a monomer fraction of >99% (n=6 independent nebulization experiments at each concentration). These results firmly underscore that the binding affinity of IN-006 was fully preserved during the process of nebulization, and no aggregates were formed.

We next evaluated IN-006 nebulized at both higher (60 mg/mL) and lower (5 and 10 mg/mL) concentrations. The APSDs of aerosols generated at 5, 10, and 60 mg/mL were very similar, as reflected by the distribution of IN-006 deposition across different stages of the NGI (Figure 2A). The MMAD for IN-006 nebulized at 5, 10, or 60 mg/mL was consistently in the range of ^~^4-5 μm, with FPF ^~^55%. The apparent difference in the MMAD compared to IN-006 nebulized at 20 and 30 mg/mL is likely due to the use of a different Innospire Go vibrating mesh nebulizer (Phillips). The binding affinity of post-nebulized IN-006 (EC_50_ ^~^5 ± 2, 5 ± 1, and 5 ± 1 ng/mL for IN-006 nebulized at 5, 10, or 60 mg/mL, respectively) was virtually identical to the affinity of pre-nebulized IN-006 (6 ± 2 ng/mL, 6 ± 2 ng/mL, and 6 ± 2 ng/mL, respectively) (Figure 2B). DLS analysis again showed no appreciable generation of aggregates with IN-006 nebulized at 5, 10, or 60 mg/mL, with >99% of IN-006 scattering present in the monomer peak (n=6 independent nebulization experiments at each concentration).

Together, these findings suggested that IN-006 can be stably nebulized at concentrations ranging from 5 to 60 mg/mL, with no loss of binding activity against SARS-CoV-2 S protein, and APSD suitable for delivery across the respiratory tract. We elected to advance our clinical formulation for IN-006 at 24 mg/mL, which supported our target daily dose of ^~^90 mg IN-006, to be administered over a span of no more than 5-7 minutes.

### GLP nebulization characterization study of IN-006

To support a formal application to regulatory authorities to initiate human studies, we next conducted nebulization characterization studies that meet Good Laboratory Practice (GLP) guidelines. In these studies, the average MMAD of IN-006 aerosols generated across 3 independent Innospire Go devices, each evaluated in triplicate, was ^~^4.6 ± 0.13 μm, with a FPF of 50 ± 1.6% (Figure 3A). The average duration to complete nebulization, with a 4.2mL fill volume, was ^~^6 ± 0.1 minutes (Figure 3A).

**Figure 3:**
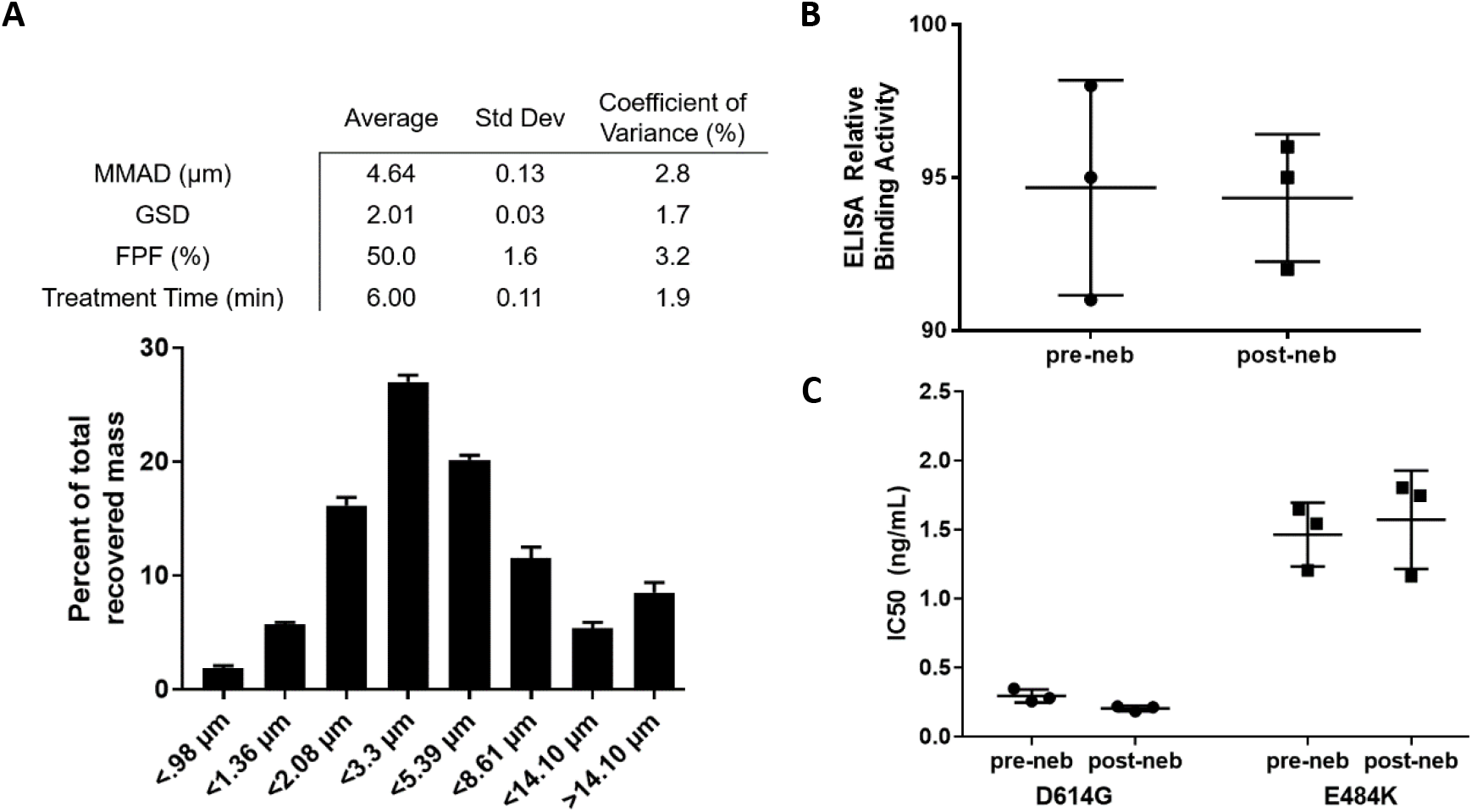
IN-006 in clinical formulation retains stable binding activity in GLP nebulization characterization study. IN-006 was formulated at 24 mg/mL and nebulized using a Phillips InnoSpire Go VMN into an NGI at 15 L/min. **A)** Aerosol particle size distribution following nebulization into NGI. Table shows summary statistics for nebulization of IN-006, including MMAD, GSD, FPF, and the treatment run-time of the nebulizer. **B)** The affinity of IN-006 was measured via Spike-binding ELISA for samples pre- and post-nebulization of IN-006 in GLP nebulization characterization studies, and there was no significant change in the relative binding affinity following nebulization (n=3 separate nebulization runs). **C)** The neutralization potency of pre- and post-nebulized IN-006 was measured against pseudotyped virus with the D614G and E484K mutations. In both assays, nebulized IN-006 provided equally strong neutralization of infection *in vitro*.

To validate the drug integrity of IN-006 following GLP nebulization, samples of nebulized IN-006 were recovered from the filters of the NGI, pooled, and evaluated against non-nebulized IN-006 stored and shipped identically as control. Comparing the relative binding affinity using a qualified RBD-coat ELISA assay, we found no difference between the pre- and post-nebulized samples with IN-006 formulated at 24 mg/mL (Figure 3B). We further confirmed the functional potency of IN-006 using pseudovirus neutralization assays. Pre- and post-nebulized IN-006 exhibited IC_50_ of 0.3 ± 0.05 ng/mL and 0.2 ± 0.02 ng/mL, respectively, against D614G pseudotyped virus, and IC_50_ of 1.5 ± 0.2 ng/mL and 1.6 ± 0.4 ng/mL, respectively, against E484K pseudotyped virus (Figure 3C). These results firmly underscore that nebulized IN-006 fully retains potent inhibition of SARS-CoV-2 infection.

To further assess the structural stability of IN-006 following nebulization and sample recovery, we conducted a rigorous series of tests to determine structural integrity, including SEC-HPLC, IEC-HPLC, and analysis for subvisible particles (Figure 4). Across all measures of drug integrity, the post-nebulized IN-006 retained excellent structural stability, with no evidence of aggregation, molecular shearing, or formation of particulate matter, compared to non-nebulized IN-006 controls. On SEC-HPLC analysis, the pre- and post-nebulized samples of IN-006 had average monomer fractions of 99 ± 0.02% and 99 ± 0.02%, respectively (Figure 4A). Similarly, CE-SDS analysis for percent intact IgG of pre- and post-nebulized IN-006 showed 98 ± 0.2% and 98 ± 0.1%, respectively (not shown). IEC-HPLC showed no differences in the mass distribution eluted in the acidic peak, main peak, or basic peak for pre- vs. post-nebulized IN-006, demonstrating stability of mAb charge variants (Figure 4B). Analysis of sub-visible particles counts showed that pre-nebulized IN-006 had 83 ± 71 particles larger than 10 μm per mL and 9 ± 8 particles larger than 25 μm per mL (Figure 4C). In comparison, post-nebulized IN-006 had 64 ± 45 particles larger than 10 μm per mL, and 1 ± 1 particles larger than 25 μm per mL (Figure 4C). The reduction in larger sub-visible particles following nebulization is likely due to a direct filtration effect by the mesh in the VMN. IEC-HPLC found equal signal for pre- and post-nebulized IN-006 in the main peak (61% and 62%, respectively), acidic group (13% and 13%), and basic group (25% and 25%). Together, these results indicate that the nebulization of IN-006 using a VMN generates a polydisperse aerosol suitable to efficiently deliver IN-006 throughout the upper and lower respiratory tract while retaining the structural and functional integrity of the molecule against SARS-CoV-2.

**Figure 4:**
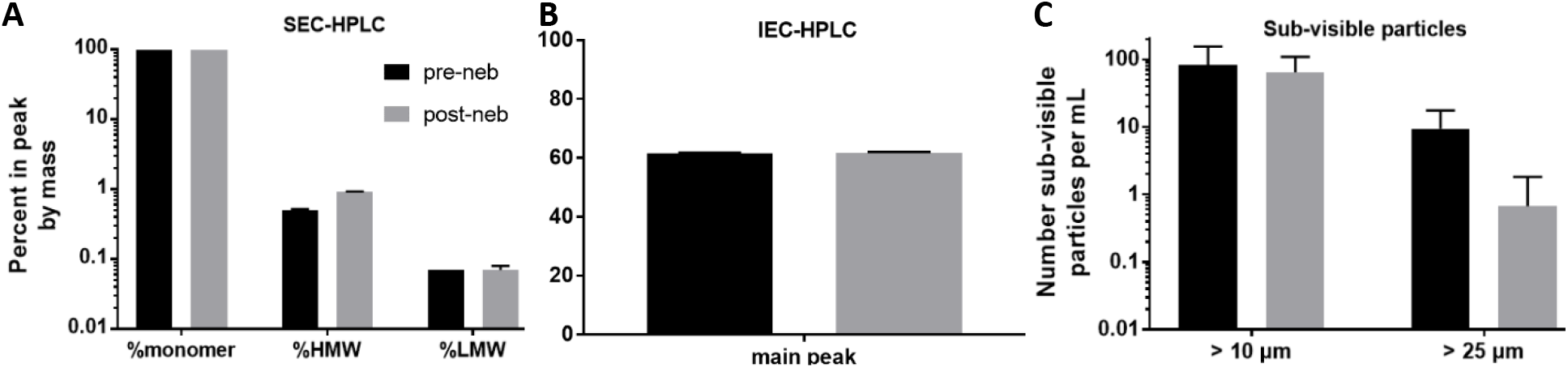
Molecular integrity of IN-006 mAb is maintained following nebulization. IN-006 was formulated at 24 mg/mL and nebulized using a Phillips InnoSpire Go VMN into an NGI at 15 L/min. Pre- and post-nebulized IN-006 was analyzed for structural integrity and impurities. **A)** SEC-HPLC analysis of the percent of mass contained in the peak representing the monomer, high molecular weight species (HMW), or low molecular weight species (LMW). **B)** IEC-HPLC analysis of the percent of mass of IN-006 eluted in the main peak. **C)** HIAC sub-visible particle analysis of the number of particles per mL that were at least 10 μm in diameter or at least 25 μm in diameter. In all physical categories assessed, nebulized IN-006 samples retained excellent physical quality attributes.

### Nebulized delivery of IN-006 achieves high concentration in the lung in vivo

Finally, we sought to determine what concentrations of IN-006 could be achieved *in vivo* in the lungs and in the systemic circulation following nebulized delivery. We treated Sprague Dawley rats with nebulized IN-006 formulated at 24 mg/mL, daily, for a period of 7 days, at a dose of either 10 or 40 mg/kg per day. From these animals, we collected broncho-alveolar lavage fluid (BALF) at 8h or 12h following the final dose (on Day 7), and serum samples at the same timepoints, from which we determined IN-006 levels using a qualified ELISA procedure. To estimate IN-006 concentrations before dilution from BALF collection, we normalized the measured IN-006 concentrations in BALF by comparing urea levels in BALF vs. urea levels in serum collected at the same time.

In the group of rats that received 10 mg/kg IN-006 daily for 7 days, the IN-006 concentrations in the lungs at 8 h and 12 h following the last dose were 183 μg/mL and 136.5 μg/mL, respectively, resulting in a calculated half-life of ^~^9.5 h, assuming steady-state, log-linear decay by 8 hours after inhaled dosing (Figure 5). In the group that received 40 mg/kg IN-006, the concentrations of IN-006 in the lungs at 8h and 12h were 1,204 μg/mL and 725 μg/mL, respectively suggesting a half-life of ^~^5.5 h in the lungs. The serum concentrations of IN-006 in the 10 mg/kg group at 8 h and 12 h were 2.3 μg/mL and 2.1 μg/mL, respectively, or ^~^60-80-fold lower than the concentrations in the lungs. The serum concentrations of IN-006 in the 40 mg/kg group at 8h and 12h after the last dose were 7.4 μg/mL and 7.1 μg/mL, respectively, or 100-160-fold lower than the concentrations in the lungs (Figure 5). Notably, while far more IN-006 was in the lungs than serum, the serum IN-006 levels were still roughly three orders of magnitude higher than the IC_50_ of IN-006 (^~^2 ng/mL, shown as dotted blue line in Figure 5). This suggests that nebulized delivery of IN-006 instantaneously achieves very high mAb levels in the lungs, but also delivers sufficient quantities of mAb into the serum to ensure effective systemic protection.

**Figure 5:**
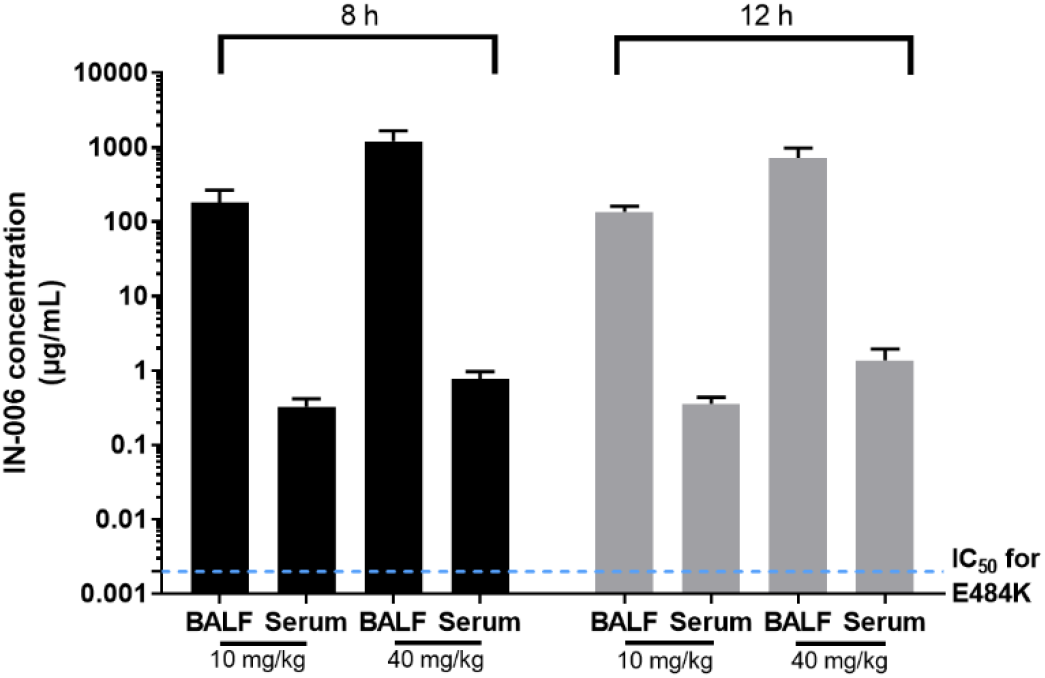
Serum and BALF concentrations of IN-006 following daily nebulized treatment in rats. Rats were treated daily with nebulized IN-006 for 7 Days. **A)** Concentrations of IN-006 in the BALF and Serum of rats at 8 h or 12 h following the final nebulized dose on Day 7, in groups that received either 10 mg/kg or 40 mg/kg. BALF IN-006 concentrations are adjusted for dilution (see Methods). Dotted blue line represents IC_50_ of IN-006 against E484K pseudovirus, ~2 ng/mL.

## Discussion

Many in the scientific community believe that COVID-19 will become endemic, despite the availability of effective vaccines [33]. This underscores the need to ensure broad availability and easy access to effective treatments that can prevent progression to severe COVID and hospitalization, particularly given the sizable population of individuals with vaccine hesitancy around the world. While highly effective mAb therapies were quickly advanced (e.g. REGEN-COV2 [34], Bamlanvimab+Etesevimab [35], Sotrovimab [36]), prior mAb treatment options have enjoyed limited adoption because of critical access issues, as well as the large doses required. First, infusions at dedicated facilities not only create a substantial burden on healthcare system and subject healthcare workers to infection risk, but also require substantial time to administer (including time for wait/registration, health check, infusion times that can last ^~^30-60 mins, and then an additional 60 mins of monitoring time post-infusion [34]). Until the creation of temporary infusion centers, many mAbs accumulated in many healthcare facilities as physicians were simply unable to administer them to enough patients [37]. While select mAbs can be given by multiple SC injections instead of IV (e.g. REGEN-COV2), that change in delivery does not appreciably ease the burden on the healthcare system, in part because patients must still spend time to register, be screened, be treated, and then still must undergo at least one hour of post-injection monitoring. To improve access, we believe it is essential to have an effective therapeutic intervention that can be used by patients in an out-patient setting immediately following a positive diagnosis using point-of-care or otherwise rapid diagnostics. Second, the systemic delivery of mAbs necessitates large mAb doses. This creates a major supply chain issue, as the sheer number of COVID-19 patients, coupled with the large mAb dose required per patient, translates to multi-metric tons of mAbs needed, which in turn creates marked constrains on global mAb manufacturing capacity.

These realities suggest an ideal treatment should not require IV, IM, or SC administrations that must be carried out by healthcare workers, or a period of monitoring following injection. Instead, the ideal treatment should be easily self-administered soon after diagnosis, allowing patients to be treated at home while still during early stages of the disease, without significant pulmonary morbidities. A safe and effective oral therapeutic clearly can meet many of the desired characteristics of convenient treatment. Two of the leading oral antivirals are Molnupiravir and Paxlovid. Molnupiravir unfortunately only provides a very modest ^~^30% reduction in relative risk of hospitalization when taken soon after outpatient diagnosis [38] (notably, substantially less efficacious than the ^~^70—87% reductions in risk provided by systemically administered mAbs [34, 35, 39]), and possess substantial safety concerns associated with mutagenesis. Paxlovid has demonstrated much greater efficacy, with an 88% reduction in relative reduction of risk of hospitalization when given to patients within 5 days of symptoms. Nevertheless, there are numerous medical contraindications for Paxlovid due to its inclusion of ritonavir, which inhibits CYP3A4, an enzyme that metabolizes many drugs (over 50% of all therapeutics [40]).

Patients are at high risk for severe COVID are more often taking multiple medications to manage other conditions, potentially limiting the use of Paxlovid as a treatment option for the very patient populations who may need an antiviral for preventing severe COVID the most. Finally, both regimens involve swallowing a substantial number of pills (40 for molnupiravir, 30 for Paxlovid) over the course of the treatment regimen, which may prove to be difficult for pediatric, geriatric, and select populations with underlying conditions [41]. While the overall prevalence of dysphagia in the Midwestern US population has been reported to be 6-9% [42], its prevalence in people over age 50 years is estimated to be closer to 15-22% [43–45], and possibly as high as 40-60% of residents in assisted living facilities and nursing homes [41, 45, 46]. For these reasons, even with the approval of Paxlovid and molnupiravir, we continue to believe inhaled treatment could address an important unmet need among COVID patients.

One of the key benefits of nebulized mAb therapy is that a much greater fraction of the overall drug dose is delivered directly to the primary site of infection and morbidity. Prior studies of antiviral mAbs for treatment of ARIs have encountered substantial roadblocks in clinical translation, likely due to inefficient and inadequate transudation of systemically administered mAb into the respiratory tract. Indeed, in a clinical study of CR6261 (an anti-influenza mAb) [21], the C_max_ in the nasal swab samples was not achieved until Day 2-3 following IV dosing, in stark contrast to peak serum concentrations of 1×10^6^ ng/ml reached within 15 minutes after infusion. More importantly, the mean peak concentration of CR6261 from nasal swabs was only ^~^600 ng/ml, or ^~^1,700-fold lower concentrations in the nasal mucosa than in plasma [21]. CR6261 is not alone in its limited distribution into the lung airways; mAbs are large molecules with generally small volumes of distribution that tend to remain in the serum and peripheral fluids in the absence of mechanisms of active transport. Previous non-human primate studies comparing the pharmacokinetics and biodistribution of systemically administered mAbs have consistently shown both slow and limited pulmonary distribution, despite achieving high concentrations in the plasma. Indeed, the concentration of mepolizumab in BALF, following IV injection, was ^~^500-fold lower than the concentration in plasma [17]. Even greater differences in BALF vs. plasma concentration were noted in biodistribution studies of motavizumab (anti-RSV mAb) in cynomolgus monkeys, where a 2,000-fold difference between BALF (^~^100 ng/mL) and plasma concentrations (^~^200,000 ng/mL) was measured 4 days following an IV dose at 30 mg/kg [18]. The poor distribution of motavizumab to the respiratory tract is likely responsible for its lack of efficacy as a treatment. In an article describing the inability of bamlanivimab to provide significant therapeutic benefit in COVID patients with severe disease, the authors hypothesized that the failure might be attributed to limited penetration into infected tissues [47]. In the case of rapidly multiplying viral infections, intervening early in the course of infection is key to effective therapy, and a few days of delay before achieving efficacious therapeutic concentrations at the site of infection may well represent the difference between clinical success and failure. For instance, treatment with oseltamivir (i.e., Tamiflu) and baloxavir marboxil (i.e., Xofluza), two oral anti-influenza antivirals, must be initiated within 48 hrs (and preferably within 24 hrs) of the emergence of symptoms in order to be efficacious [48]. Since IN-006 can achieve pulmonary C_max_ virtually instantaneously, we believe it is exceptionally suited as an early intervention to prevent progression to severe COVID.

The frequent reports of mAb aggregation following nebulization [22] have led many in the field to believe fully human mAbs are too large or too unstable to be nebulized, and that smaller protein binders such as nanobodies (camelid antibodies that consist of only heavy chains, without light chains and without effector functions) are more suitable for nebulized delivery. In sharp contrast to this prevailing dogma, we showed here that nebulization of IN-006, a fully human IgG1 mAb, did not result in any appreciable increase in aggregation or fragmentation, a loss of binding/neutralization activity, or other impacts on physical integrity (e.g., monomer content). Although mAbs and nanobodies would both be expected to provide potent neutralization, the presence of an Fc domain on the full mAb confers additional effector functions, including opsonization, ADCC and ADC, and muco-trapping.

Further, since nanobodies are camelid in origin, there may be substantial immunogenicity, as previously reported for inhaled nanobodies for RSV treatment. For these reasons, we believe nebulized treatments using full length human IgG1 mAbs may confer additional benefits over nebulized therapies based on nanobodies or Fabs.

The potential muco-trapping effector function of IgG – to physically trap pathogens in mucus – has only been appreciated recently. An inherent assumption by the field has been that, to trap a pathogen, Ab must bind tightly to mucins. However, many investigators reported seemingly negligible affinity of IgG to mucins [49–53]; for instance, the diffusion rates of IgG in human cervical mucus are slowed only ^~^10% vs. their rates in buffer, implying that 90% of the time an IgG is simply not bound to mucins [54]. This led most researchers to conclude that Ab have no meaningful function in mucus besides neutralization. Instead, our discovery of the muco-trapping potential of IgGs is based on multiple weak and transient bonds between IgGs and mucins, and highlights two pivotal concepts: First, many IgGs can bind to the surface of a pathogen, and the resulting array of IgGs can generate high binding avidity to mucins (analogy: a Velcro® patch can tightly bind two surfaces despite individually weak hooks). Second, IgG must possess a narrow range of weak and transient affinity to quickly accumulate on the invading pathogen surface while minimizing the number of pathogen-bound IgG needed to trap the pathogen [55–58]; mAbs that bind too tightly to mucins would not be able to travel through mucus to bind pathogens. We have previously shown that IgG possessing suitable N-glycans on IgG-Fc is capable of immobilizing viruses in various mucus secretions, resulting in rapid clearance from the respiratory tract [59] and can provide effective protection against vaginal Herpes transmission [54, 57]. In good agreement with our previous findings, we showed here that IN-006 was able to effectively immobilize SARS-CoV-2 in AM. Trapped virions are unable to diffuse through AM to infect cells and are expected to be quickly eliminated from the respiratory tract by natural muco-ciliary or cough-driven mucus clearance mechanisms for sterilization in the low pH gastric environments. Thus, mAbs capable of this muco-trapping effector function provide a mechanism to *physically* remove viruses from infected airways.

The molecular target for SARS-CoV-2, ACE-2, is expressed on cells throughout the entirety of the respiratory tract, in a gradient with greatest expression in the upper airways and progressively decreased expression in the lower respiratory tract [60]. Although SARS-CoV-2 infections most likely initiate in the nasal turbinates, it is impossible to predict, for each individual patient, whether the infection is still strictly localized in the nasal passage at the time of a confirmed positive diagnosis, or whether some of the virions may have already disseminated to the lower respiratory tract. Thus, for a topical therapeutic to be maximally effective for patients in all stages of early infection, we believe there is a need to deliver the drug to all parts of the respiratory tract, rather than target only the lungs (e.g., with the smaller particles typically generated by dry powder inhalers) or target only the upper airways (e.g., with the application of nasal drops). We believe nebulization, by generating diverse droplet sizes that enable simultaneous delivery to all parts of the respiratory tract, is uniquely suited for inhaled delivery of antivirals.

## Acknowledgments

This work was financially supported in part by the United States Army Medical Research Institute of Infectious Disease (USAMRIID) through Medical Technology Enterprise Consortium (MTEC) award W81XWH-20-9-0008, by the North Carolina Policy Collaboratory (S.K.L.), and the David and Lucile Packard Foundation (2013-39274; S.K.L). The content is solely the responsibility of the authors and does not necessarily represent the official views of the funding organizations.

## Declaration

MM, ZR, and JH are employees of Inhalon Biopharma/Mucommune, and hold shares in Inhalon Biopharma. HK, ML, CK, and KK are employees of Celltrion, Inc. SKL is founder of Mucommune, LLC and currently serves as its interim CEO. SKL is also founder of Inhalon Biopharma, Inc, and currently serves as its CSO as well as on its Board of Director and Scientific Advisory Board. S.K.L has equity interests in both Mucommune and Inhalon Biopharma; S.K.L‘s relationships with Mucommune and Inhalon are subject to certain restrictions under University policy. The terms of these arrangements are managed by UNC-CH in accordance with its conflict of interest policies.

## Materials and Methods

<u>*Production of*</u> fluorescent VLPs were made by co-transfection of pGAG-mCherry plasmid (kind gift from Gummuluru lab) and Cov2 S protein plasmid in a 1:1 ratio. Non-replicating lentivirus pseudotyped with SARS-CoV-2 UK spike protein was created using the following plasmids, in a 1:1:1:2 ratio: pMDLg/pRRE, pRSV-REV, SARS-CoV-2 UK Spike, and pLL7 GFP. The plasmids were transfected into LVMaxx using the LVMaxx Transfection kit. Each VLP was made in 60mL cultures, and harvested after 48 hours. The VLPs were purified using 25% Sucrose (in 25mM HEPES/130mM NaCl) cushion spin protocol. 3mL of 25% sucrose solution was add to each Beckman Coulter ultracentrifuge tube, which then had 7mL of cell culture supernatant gently layered on top. The tubes were then spun at 36,000rpm for 2.5 hours at 4°C. The sucrose/supernatant was then aspirated off, and 20μL of 10% Sucrose solution was placed on top of the VLP pellet. After 24 hours at 4°C, the VLPs were then aliquoted and stored at −80°C.

### Multiple particle tracking for quantifying mobility of SARS-CoV-2 virus like particles in AM treated with IN-006

Fresh, undiluted human AM were collected from extubated and otherwise would-be-discarded endotracheal tubes, as previously described [59], via a non-human subjects research designed protocol approved by the University of North Carolina at Chapel Hill. All AM were kept on ice, treated with protease inhibitors, and used within 24-72 hrs of collection. Multiple particle tracking analysis of fluorescent SARS-CoV-2 VLPs in human AM was performed as described in [59]. Briefly, solutions of fluorescent VLPs and IN-006 were added to ^~^10 μL of fresh, undiluted airway mucus in custom-made glass chambers. The samples were then incubated at 37 °C for ^~^30 mins before microscopy. The same AM was used for all conditions to allow direct comparison among samples. Videos of VLPs diffusing in AM were recorded with MetaMorph software (Molecular Devices, Sunnyvale, CA) at a temporal resolution of 66.7 ms. Videos were analyzed using NetTracker from AI Tracking Solutions to convert video raw data to particle traces. Time-averaged mean-squared displacements (MSDs) and effective diffusivity were calculated by transforming particle centroid coordinates were transformed into time MSDs with the formula <Δ*r*^2^(τ)> = [*x*(*t* + τ) – *x*(*t*)]^2^ + [*y*(*t* + τ) – *y*(*t*)]^2^, where τ = time scale or time lag.

### Non-GLP Nebulization study

To test the feasibility of nebulizing IN-006 across a range of concentrations, IN-006 was formulated at various concentrations from 5-60 mg/mL and nebulized using an InnoSpire Go vibrating mesh nebulizer. Generally, USP <1601> was adhered to for generation of data and calculation of mass median aerodynamic diameter (MMAD) and geometric standard deviation (GSD). As discussed in this report, fine particle fraction refers to particles collected on stages 4 and smaller of the NGI (< 5.39 μm at 15 L/min). Briefly, the Next Generation Impactor (NGI; MSP Corp, MN, USA) and collection stages were pre-cooled to 4 °C for at least 90 minutes prior to experiments. The nebulizer was loaded with enough mAb solution to ensure replicates could be performed sequentially, while avoiding sputtering (i.e., remaining above the manufacturer’s minimum recommended volume). A custom silicone mouthpiece molded to the nebulizer/NGI inlet interface was used to affix the nebulizer to the inlet with a tight seal. A solenoid in line with the NGI and vacuum (set to 15 L/min) was used to collect sufficient nebulized mAb at a given concentration. The nebulizer was actuated, and the solenoid was switched on to begin collection. Following nebulization, the vacuum and nebulizer were switched off and the NGI stages and inlet were removed. Quickly, the next set of stages and inlet were swapped in to perform a second replicate nebulization. To collect deposit mAb, stages were rinsed with 5 mL of the formulation buffer, matching the buffer of nebulized material, and assayed for mAb mass deposition at A280. APSDs were plotted as cumulative percentage of drug mass undersize against aerodynamic stage cut-off diameter for IN-006 on a log-probability scale. The MMAD, GSD, and FPF were determined from this data.

In assessing APSD, the median mass aerodynamic diameter (MMAD) was defined as the aerosol diameter cutoff at which 50% of the mass of drug was in larger aerosols and 50% was in smaller particles. The fine particle fraction (FPF) is calculated as the mass of drug contained in particles smaller than ^~^5 μm divided by the total emitted dose to roughly estimate the fraction of nebulized therapeutic that would be delivered to the lower airways.

### ELISA for determining binding affinity pre- and post-nebulization in non-GLP studies (Figure 2)

ELISA binding assays were performed using 96 well half-area plates (Fisher Scientific, Costar 3690) coated with 0.5 μg/mL of S protein and incubated overnight at 4°C. Plates were washed with PBS with 1:2000 Tween 20. ELISA plates were blocked the following day with 5% (w/v) milk in PBS with Tween 20 at a 1:2,000 dilution at room temperature for one hour. Samples and standard curves were diluted in 1% (w/v) milk in PBS with Tween 20 at a 1:10,000 dilution. Samples were incubated at room temperature for 1hr. Plates were washed with PBS containing Tween 20 at a 1:2000 dilution three times. The detection antibody was a peroxidase-conjugated goat anti-human IgG Fc antibody (Rockland 709-1317), used at a 1:5000 dilution in 1% milk in PBS, and was incubated at room temperature for 1hr. The solution was then discarded and washed three times. Plates were developed with TMB solution, and development was stopped by adding 2N HCl. The absorbance at 450 nm (signal) and 570 nm (background) was then measured with a microplate photodetector (Fisher Scientific).

### Dynamic Light Scattering for determining aggregates in non-GLP study

To detect the presence of aggregates following nebulization, post-nebulization samples were collected from the NGI and measured via dynamic light scattering (Zetasizer, Malvern Instruments, particle quantitation limit 0.3 nm – 10 μm). Post-nebulized IN-006 samples were added to a cuvette at the collected concentration (^~^1 mg/mL after dilution during sample recovery), and particle size polydispersity and average diameters were determined using volume-weighted analyses.

### GLP Nebulization study and characterization

Three InnoSpire Go devices were tested in triplicate into a cascade impactor (Copley NGI), operated with an extraction flow of 15 ± 0.5 L/min and temperature of 5 ± 2 ºC. Each device was loaded with a total of 4.2 mL of IN-006. Gravimetric weights were recorded to enable full mass balance calculations. Devices were operated for 90 sec into the NGI. At the end of the run, samples were collected from all stages of the NGI and analyzed by UV microplate reader (A280). This nebulization method was used to determine delivered dose, APSD, and the collected samples were used to assess drug integrity following nebulization (e.g., ELISA, neutralization assay, and HPLC).

#### SARS-CoV-2 RBD binding ELISA (for GLP nebulization characterization study; Figure 3B)

Post-nebulized samples were serially diluted (1,200 ng/mL – 0.00239 ng/mL, 10 points) and added to a 96 well microplate previously coated with 0.05 μg/mL of SARS-CoV-2 RBD manufactured by Celltrion. The bounded material was detected using anti-human IgG Fc-HRP conjugate. The signal was measured after TMB (3, 3’, 5, 5’-tetramethylbenzidine) treatment. The optical density values were fitted using 4-PL logistic analysis and the relative binding activity of sample to SARS-CoV-2 RBD was determined from comparison of the EC50 value of samples and that of CT-P59 in-house reference standard by PLA software.

#### Pseudovirus Neutralization

SARS-CoV-2 spike mutant pseudovirus was produced by transfection into HEK-293T with plasmid mixture such as third-generation Lentiviral packaging vectors pMDLg/pRRE, pRSV-Rev, pCDH-CMV-Nluc-copGFP-Puro, a dual reporter vector expressing luciferase and GFP, and the pCMV3-SARS-CoV-2 spike plasmid expressing the mutant SARS-CoV-2 spike prepared through each site-direct mutagenesis. Mixture of serially diluted antibodies at 3-fold ratio from 100 to 0.005 ng/ml and diluted pseudovirus to a predetermined number of copies was incubated for 1 hour and was added to stably expressing human ACE-2 HEK293T cells which are seeded into 96-well plate the day before the test. After 24 hours of incubation, the medium was replaced with fresh medium. At 72 hours post-infection, pseudovirus neutralizing activity was measured using Passive lysis buffer (Promega) and Nano-Glo Luciferase Assay Reagent (Promega) according to the manufacturer’s manual. Inhibition% was calculated based on the average value of RLU (Relative Luminescence Units) of virus only control. Finally, IC_50_ value was calculated using a nonlinear regression model by GraphPad Prism 5.0.

#### SEC-HPLC

Size exclusion chromatography was performed to evaluate aggregates, monomer and fragments ratio under non-denaturing conditions for pre- and post-nebulization samples. This assay was performed by Waters HPLC Alliance system on a TOSOH TSKgel G3000SWXL column (7.8 mm*300 mm) using aqueous buffered mobile phase at ambient temperature. The isocratic elution profile at a constant flow rate 1.0 mL/min was monitored using UV detection at 214 nm.

#### IEC-HPLC

Ion exchange chromatography was performed to evaluate the distribution of charge variants pre- and post-nebulization samples using cation exchange chromatography. The Waters HPLC Alliance system was equipped with a BioPro IEX SF analytical column (4.6*100 mm) at ambient temperature. Gradient NaCl elution was performed at a constant flow rate of 0.8 mL/min and UV signals were obtained at 280 nm.

#### Sub-visible Particles using Light-obscuration (HIAC)

The number of sub-visible particles (SVPs) in pre- and post-nebulization samples were measured by light obscuration method using the HIAC liquid particle counting system. Analysis was performed by HACH ULTRA analytics, HIAC 9703 liquid particle counter equipped with 1.0 mL syringe and small bore probe at ambient temperature. Processing was performed using Pharm spec software.

### Rat nebulization study

To determine the concentrations of IN-006 achievable *in vivo* following inhaled dosing, we treated male and female Sprague Dawley Rats (age ^~^9-10 weeks at start of treatment) with nebulized IN-006 daily in an inhalation chamber, for 7 days. On Day 7, BALF and serum samples were collected at 8 h and 12 h following the last dose of IN-006. Stability of nebulized IN-006 in this exposure system was verified by collecting a sample of aerosolized material at one of the exposure ports and validating maintenance of anti-SARS-CoV-2 binding activity (via RBD-coated ELISA with EC_50_ within 20% of pre-nebulization control). Exposure at a dosing level of 10 mg/kg per day or 40 mg/kg per day was achieved by first determining the rate of IN-006 output in the exposure chamber through sampling aerosolized material using a glass fiber filter (Whatman® glass microfiber filters, Grade GF/A circles). Then, we chose a corresponding duration of exposure to IN-006 nebulized at 24 mg/mL to achieve 10 mg/kg or 40 mg/kg per day (i.e., we changed duration of exposure to IN-006 treatment, and did not change the formulation concentration of IN-006 to achieve different dosages).

### Urea Measurement for determination of BALF dilution factor

We sought to determine the extent to which the BALF samples were diluted during collection (rinsing of the lungs with saline). During this collection process, the airway fluid is inherently diluted, artificially decreasing the apparent concentration of therapeutics in the recovered BALF. We therefore measured the concentration of urea in the BALF samples and in the serum samples to determine the average extent of dilution (prior to dilution, urea concentrations are equal in the lung fluid and in the serum, allowing for the calculation of a dilution factor [61]). On average, the urea concentrations in the BALF were 31.4-fold lower than those measured in the serum, suggesting an approximate dilution factor of 31.4× during BALF collection. The concentrations of urea in BALF and serum were measured using a commercial kit (Abcam: ab83362). The kit allows quantification of urea in a variety of biological samples such as serum, BALF, urine, etc. Samples of rat serum or BALF were diluted in a range of 1:50-1:200 for serum, or 1:10-1:20 for BALF, and the apparent concentrations of urea were determined through measurement of absorbance at 570nm and comparison to a linear standard curve with standards between 0.5 nmol urea per well to 5 nmol urea per well. Fold-dilution of BALF was calculated as the concentration of urea in serum divided by the concentration of urea in BALF. Example: serum urea concentration (4 mM), BALF urea concentration (0.1 mM): fold-dilution of BALF (4 mM / 0.1 mM = 40× dilution factor).

---

**Supp Table 1.** Information about donors who provided airway mucus samples. All donors had no upper respiratory tract infection and no pulmonary disease at the time of collection.

| Age | Gender | Smoker or Exposure to smoker | Race/Ethnicity |
| --- | --- | --- | --- |
| 60 | F | N | Caucasian |
| 13 | F | N | Caucasian |
| 65 | M | N | Caucasian |
| 56 | M | Y | Hispanic |
| 54 | F | N | Caucasian |
| 31 | M | N | African American |
| 21 | M | N | Caucasian |
| 64 | F | N | Caucasian |
| 55 | M | N | Hispanic |
| 64 | F | N | African American |

